# Task-dependent pupillary responses to glossiness and attractiveness judgments

**DOI:** 10.1101/2025.11.05.686874

**Authors:** Hideki Tamura, Shigeki Nakauchi, Tetsuto Minami

## Abstract

Human pupillary responses are influenced not only by low-level visual properties but also by cognitive and affective factors related to task demands. However, the temporal dynamics of how different evaluative tasks modulate pupil size remain poorly understood. In this study, we investigated how pupillary responses vary when observers evaluate the same set of object images for either glossiness or attractiveness. These two perceptual attributes were selected to exemplify distinct cognitive demands: one rooted in surface-level visual analysis and the other involving emotional valuation. The stimuli consisted of 60 grayscale object photographs selected from the THINGS database, representing common real-world items without social or facial content. Low-level image statistics were controlled using histogram matching to equalize luminance and contrast across all stimuli. Participants viewed each image for 3000 ms while maintaining central fixation and rated either its glossiness or attractiveness on a 7-point scale in separate task blocks. Ground-average waveforms revealed task-dependent modulations of pupil size across rating levels, consistent with prior reports that pupillary responses vary with evaluative context even for identical stimuli. Using temporal principal component analysis and generalized additive modeling, we found that higher glossiness ratings were associated with greater pupil constriction at early, light-reflex-like latencies, whereas higher attractiveness ratings elicited greater pupil dilation at later time points. These findings suggest that distinct temporal profiles of pupil size reflect task-specific processing demands, potentially aligning with the notion that visual and affective evaluations may unfold in temporally distinct stages, as suggested in prior theoretical models. Our results underscore the value of pupillometry as a non-invasive tool for dissociating task-dependent perceptual processes, with potential applications in cognitive neuroscience and affective computing.

**Impact statement:** This study reveals that pupil size reflects not only what we see, but also how we evaluate it. By showing distinct temporal signatures of pupillary responses for glossiness versus attractiveness judgments of the same images, our findings suggest temporally dissociable stages of sensory and affective processing. These results highlight the value of pupillometry as a non-invasive index of task-dependent cognitive evaluation.

## 1. Introduction

Humans perceive a rich array of material qualities (shitsukan, a Japanese term referring to perceived material attributes such as texture, gloss, or softness) through vision (Adelson, 2001; Anderson, 2011; Anderson & Marlow, 2022; Fleming, 2014, 2017; Komatsu & Goda, 2018). These perceptual qualities span a spectrum from low-level, physically grounded attributes—such as glossiness, surface texture, and material composition—to high-level, affect-laden impressions, including preferences, aversions, and emotional valence (Komatsu & Goda, 2018). The former qualities are typically processed automatically on the basis of physical cues, where as the latter qualities involve abstract evaluations shaped by emotional and contextual factors. Komatsu & Goda (2018) proposed a hierarchical model of material perception, positing that such perceptual attributes are processed along a continuum in the visual system, from low-level sensory features to higher-order cognitive evaluations (Komatsu & Goda, 2018). For instance, glossiness is primarily processed within the ventral visual pathway (Baba et al., 2021; Komatsu et al., 2021; Nishio et al., 2012, 2014; Okazawa et al., 2012), whereas aesthetic impressions such as attractiveness are thought to engage higher-order regions, including the prefrontal cortex and reward-related areas (Chatterjee et al., 2009; Cinzia & Vittorio, 2009; Kawabata & Zeki, 2004; Kirk, 2008; O’Doherty et al., 2003). In this framework, gloss perception is driven largely by low-level features such as luminance, orientation, and shape, whereas attractiveness perception builds upon these visual cues to support affective and value-based judgments. This hierarchical processing is theoretically supported by psychophysical findings, particularly those involving temporal indicators such as stimulus exposure duration and response latency (e.g., Nagai et al., 2015). However, its physiological correlates—especially those captured by psychophysiological indices—remain largely unexplored.

One of the key psychophysiological indicators of internal cognitive and emotional states is pupil size. While pupil diameter is known primarily for its reflexive adjustment to changes in luminance, considerable research has demonstrated that it is also highly sensitive to attentional processes (Binda et al., 2013a; Bombeke et al., 2016; Mathôt et al., 2013; Naber et al., 2013) and emotional arousal (Bradley et al., 2008; Kuraguchi & Kanari, 2020, 2021; Laeng et al., 2013; Oliva & Anikin, 2018). These cognitively driven modulations of the pupil, referred to as psychosensory responses (Ebitz & Moore, 2019; Mathôt, 2018), are believed to involve subcortical structures such as the superior colliculus and locus coeruleus, which regulate early attentional orienting and arousal-related pupil dilation, respectively.

In contrast, pupil constriction in response to perceived brightness—even in the absence of changes in physical luminance—may reflect the engagement of cortical pathways involved in interpreting surface appearance, although the underlying mechanisms remain to be fully elucidated. For instance, Tamura et al. (Tamura et al., 2024) reported a significant association between the perceived glossiness of everyday objects and pupil constriction, particularly in the context of the pupillary light reflex. This finding suggests that even relatively low-level perceptual qualities, such as glossiness, can modulate the pupillary system depending on the degree of subjective perception. Such psychosensory responses involving pupil constriction have also been documented in response to visual illusions such as the glare effect, where perceived brightness is exaggerated beyond physical stimulus properties (Kinzuka et al., 2021; Suzuki, Minami, Laeng, et al., 2019; Suzuki, Minami, & Nakauchi, 2019; Zavagno et al., 2017). However, although prior studies have revealed that subjective impressions of visual materials can modulate pupil responses, it remains unclear whether and how different types of perceptual evaluations—such as lower-level judgments of surface appearance and higher-level assessments of affective value—engage distinct pupillary mechanisms. While previous work has highlighted the multifactorial nature of pupil responses—including sensitivity to luminance, attention, and affect—relatively little is known about whether different evaluative dimensions are processed at distinct temporal stages. Theoretical models of material perception suggest a hierarchical structure in which low-level visual cues (e.g., glossiness) are processed earlier than more affective or semantic dimensions (e.g., attractiveness) (Komatsu & Goda, 2018). However, this hypothesis has rarely been tested using time-resolved physiological indices such as pupillometry.

To address this research gap, the present study focused on two types of texture evaluation tasks that are assumed to differ in their level of processing: perceived glossiness, which is primarily based on physical and visual cues (i.e., a lower-level texture quality), and perceived attractiveness, which involves affective and value-based judgments (i.e., a higher-level texture quality). By comparing pupil responses to these distinct evaluative tasks while the same visual stimuli were presented, we aimed to clarify how the level of abstraction or cognitive demand in material perception is reflected in physiological signals. This approach may offer empirical insights into how perceptual attributes of different abstraction levels are temporally reflected in physiological signals—an idea that aligns with theoretical accounts of material perception such as that proposed by Komatsu & Goda (2018). In particular, we hypothesize that pupil responses may differ in their temporal characteristics, with glossiness-related responses occurring earlier and attractiveness-related responses emerging later. If such temporal dissociation is observed even with identical stimuli and if the magnitude of pupil change depends systematically on subjective ratings in each task, distinct cognitive processes may be associated with different evaluative dimensions, resulting in distinct temporal patterns in pupil dynamics. Importantly, our goal is not to directly test the hierarchical model itself but rather to describe and characterize task-specific differences in pupil response and to explore their neurocognitive implications within that broader theoretical framework.

To test this hypothesis, we conducted a psychophysical experiment in which participants evaluated the same set of visual stimuli under two perceptual tasks—rating glossiness and attractiveness—to examine how the temporal profile of pupillary responses varies as a function of task demands. Prior research has demonstrated that even when visual input is held constant, differences in evaluative goals or cognitive framing can lead to marked changes in pupil dynamics, including task-dependent temporal shifts. For instance, Liao et al. (2021) presented participants with the same facial images but instructed them to either provide an attractiveness rating or perform a shape discrimination task (Liao et al., 2021). Their findings revealed a transient pupillary constriction that was particularly pronounced during attractiveness judgments. Interestingly, a similar constriction was observed during shape discrimination, suggesting that attractiveness-related processing may influence pupil size even in the absence of explicit evaluation. Similarly, Santos et al. (2023) showed participants images of hands holding objects while the evaluative context was manipulated: participants were told that the hands were either “contaminated and infectious” or “clean” (Santos et al., 2023). Under the contamination framing, participants reported higher levels of disgust and exhibited stronger pupillary constriction. These studies underscore that task context and evaluative instructions can modulate pupillary responses, highlighting the sensitivity of this physiological measure to cognitive and affective appraisals—even in the absence of changes in visual input.

In this study, we conceptualize glossiness as a lower-level perceptual attribute grounded in visual cues and attractiveness as a higher-level perceptual construct involving emotional and valuation processes. On the basis of this distinction, we hypothesized that pupil responses would differ in both direction and timing depending on the evaluative task: glossiness judgments would trigger relatively early pupillary constriction, consistent with light-reflex mechanisms; in contrast, attractiveness judgments were expected to elicit delayed pupil dilation, which is indicative of emotionally driven arousal responses. To test this hypothesis, we employed two complementary analytic approaches that capture both localized and global temporal aspects of pupillary dynamics. First, we applied temporal principal component analysis (tPCA) (e.g., Scharf et al., 2022) to extract the dominant time-varying components of pupil responses and examined how their peak amplitude and latency varied as a function of task and subjective ratings. This approach allows for a temporally resolved characterization of how the evaluative context shapes pupil dynamics. For example, Blini et al. (2024) demonstrated that tPCA can identify consistent latent structures—termed “pupil manifolds”—across different tasks and stimuli, suggesting that pupil responses may be shaped by intrinsic physiological constraints (Blini et al., 2024).

Building on this insight, we tested whether distinct pupillary components would be obtained when identical stimuli were evaluated under different task instructions. Second, we used generalized additive models (GAMs) to model nonlinear interactions among task, rating, and time, thereby capturing the full temporal trajectory of pupil size. This method allows us to detect when and how the pupil response diverges across conditions while accounting for individual variability via random effects (Rij et al., 2019). In particular, GAMs enabled us to examine not only differences in response magnitude but also the onset and duration of dilation, which are critical for testing our prediction that attractiveness judgments would be accompanied by delayed, sustained pupil dilation. Even with constant visual inputs, these two analytic approaches enabled us to reveal how distinct types of perceptual evaluation modulate both the timing and structural dynamics of pupillary responses. By directly comparing pupil responses to glossiness and attractiveness judgments of the same object images, we aimed to test whether the evaluative context modulates the temporal dynamics of pupil size. Such task-dependent modulation provides indirect support for the hypothesis that material perception occurs in distinct stages, which is consistent with—but does not directly test—the hierarchical framework.

## 2. Methods

### 2.1 Participants

Twenty-four naïve observers participated in the experiments. The sample size was determined on the basis of a previous study that investigated the relationship between perceived glossiness and pupillary responses (Tamura et al., 2024). Participants had normal or corrected-to-normal visual acuity, and the mean age of the participants was 22.4 ± 1.0 years. All experimental procedures were approved by the Institutional Review Board of Toyohashi University of Technology. The study was conducted in accordance with ethical standards consistent with the Declaration of Helsinki. Written informed consent for the publication of participants’ information was obtained prior to participation.

### 2.2 Apparatus

The experiment took place in a darkened booth under low ambient illumination (approximately 60 lx). Visual stimuli were presented on a 27-inch LCD monitor (ColorEdge 27 CS2731, EIZO) with a resolution of 1920 × 1080 pixels and a refresh rate of 60 Hz, which was calibrated using a SpyderX Elite system (ImageVISION). The monitor was luminance-calibrated such that the white point reached 120 cd/m². Participants sat in the booth with their heads stabilized by a chin rest, ensuring a constant viewing distance of 86 cm. Pupil diameter was recorded using an EyeLink Portable Duo eye tracker (SR Research) at a sampling rate of 500 Hz, with a five-point calibration procedure conducted before each session. Participants provided their responses using a numeric keypad. The stimulus presentation was controlled via MATLAB with the Psychtoolbox 3.0 extensions (Brainard, 1997; Kleiner et al., 2007; Pelli, 1997).

### 2.3 Stimuli

A total of 60 object images were selected from the THINGS database (Hebart et al., 2019, 2020), which contains photographs of real-world items commonly encountered in daily life. These images were identical to those employed in our previous study (Tamura et al., 2024). To curate a stimulus set that broadly represents general objects while minimizing potential confounding factors related to glossiness and attractiveness, we implemented a multistage selection procedure prior to the main experiment. First, a naïve individual who did not participate in the main study categorized 1,854 object names from the THINGS database into three broad semantic categories: objects, creatures, and food.

Of these, 209 items labeled as creatures and 284 labeled as food were excluded, as we focused on objects that are typically associated with surface materials likely to vary in glossiness and attractiveness, which aligned with our experimental goals. Next, the perceived glossiness and luxury of the remaining 1,361 object names were evaluated by the same individual, with luxury used as a proxy for attractiveness. This step was performed to encourage grouping on the basis of semantic and functional characteristics of the objects rather than on purely affective or aesthetic impressions. Although luxury and attractiveness are conceptually distinct, they are assumed to be substantially correlated, making luxury a reasonable substitute in this context. On the basis of this preliminary categorization, 60 object images were selected such that the distributions of glossiness and attractiveness were approximately uniform across the set. This strategy helped prevent unintentional clustering in either dimension and allowed the two rating axes to be manipulated independently during the main task. Each selected image depicted a single, clearly visible object with a sufficiently large surface area available for visual evaluation.

Each of the 60 selected images was drawn from a distinct category defined in the THINGS database. The images were resized to 512 × 512 pixels, converted to grayscale, and cropped to remove the background while preserving the main object. To control for low-level visual features, we applied histogram matching using the SHINE toolbox (Willenbockel et al., 2010), aligning the luminance histograms of all the images to the average across the set. This procedure equalized key image statistics, resulting in the following matched values: mean luminance = 27.24 ± 0.68 cd/m², variance = 795.00 ± 38.78, skewness = 1.25 ± 0.06, and kurtosis = 3.71 ± 0.20.

### 2.4 Procedure

The experimental procedure is illustrated in Figure 1. Each trial began with a 1000-ms interstimulus interval (ISI), followed by the presentation of a black fixation cross (0.6 × 0.6 degrees, 0.15 cd/m²) on a gray background (9.19 cd/m²) for 1000 ms. A stimulus image then appeared at the center of the screen, together with the fixation cross, and remained visible for 3000 ms. Each stimulus depicted a single object positioned within a 6 × 6 degree region of the visual field. Participants were instructed to rate the object using a 7-point Likert scale (1 = low, 7 = high). Two types of rating tasks were employed: one assessing subjective attractiveness and the other assessing subjective glossiness. In both tasks, participants were instructed to base their ratings on their personal impressions rather than on any objective criteria. Responses were entered using a numeric keypad. The next trial began immediately after a response was made. Participants were asked to maintain central fixation whenever the fixation cross was visible, and to blink only during the ISI period when the screen was blank. Each participant completed a total of 240 trials, comprising 60 images × 2 rating tasks × 2 repetitions. The two rating tasks were administered in separate blocks of 120 trials each. The order of these blocks was counterbalanced across participants, and the trial order within each block was randomized. A five-point calibration of the eye-tracking device was performed at the beginning of each block. Participants were allowed to take self-paced breaks between blocks.

**Figure 1.**
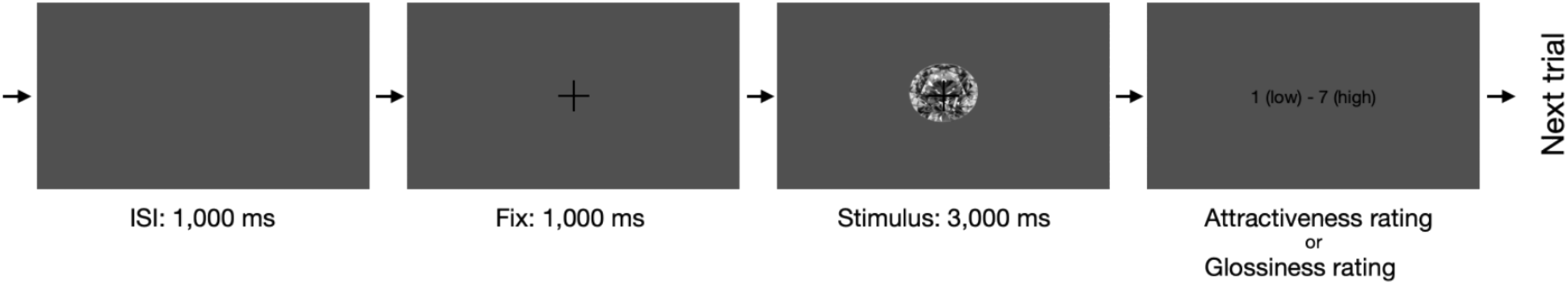
Procedure. Schematic sequence of a single trial in the experiment. Note that the relative sizes of the fixation point and stimulus to the screen are not to scale and differ from those used in the actual experiment.

### 2.5 Data analysis

Behavioral and pupillometric data were analyzed using R (version 4.3.3) and MATLAB R2023a. We computed the mean attractiveness and glossiness ratings for each image on the basis of participants’ responses. Only trials retained for the pupillometric analysis were included; those excluded because of signal artifacts or missing data were also omitted from the behavioral analysis to ensure consistency.

Pupil diameter was analyzed from the onset of the fixation cross to the end of stimulus presentation. Preprocessing was performed following procedures established by Nakakoga et al. (2021) and Tamura et al. (2024), with data from the right eye used in the analysis. Eye blinks were identified as time intervals in which the pupil velocity exceeded 0.011 mm/ms. These intervals, along with 10-ms buffers on either side (i.e., a total of 20 ms), were excluded. The missing segments were interpolated using a shape-preserving piecewise cubic spline function. Entire trials were excluded from analysis if they contained a blink lasting longer than 1 s, if more than 30% of the data points were missing, or if the pupil signal was absent at either the beginning or end of the trial. Participants with more than 50% of trials rejected were selected to be excluded on the basis of preestablished criteria; however, no participant met this threshold. Consequently, data from all 24 participants were retained for analysis (mean rejection rate: 12.07% ± 13.98%). Baseline correction was performed by subtracting the average pupil diameter measured during the 200 ms preceding stimulus onset. To reduce high-frequency noise, the baseline-corrected data were smoothed using a 10-ms moving average filter.

Pupillary responses were first averaged across trials for each task and rating level to obtain grand-average waveforms. In addition, tPCA was conducted following the method proposed by Scharf et al. (Scharf et al., 2022). The resulting factor loadings were multiplied by the original pupil data to compute the pupil component scores using an approach analogous to that described by Wetzel et al. (Wetzel et al., 2020). To further examine the continuous effects of task on pupillary dynamics, GAMs were applied to the component scores for each factor.

## 3. Results

### 3.1 Behavioral results

Figure 2a shows histograms of the mean attractiveness and glossiness ratings for each image. The attractiveness ratings followed a slightly broad unimodal distribution, whereas the glossiness ratings exhibited a bimodal distribution with peaks at approximately 3 and 5. The average attractiveness and glossiness ratings across the 60 images were 3.97 ± 1.03 and 3.90 ± 1.24, respectively. The two ratings were positively correlated (𝑟 = .42, 95% CI [.19, .61], 𝑡(58) = 3.53, 𝑝 < .001). Figure 2b illustrates the distribution of ratings for each stimulus in the two-dimensional space of attractiveness and glossiness. Objects rated high on both dimensions included gemstones (stimulus ID = 18) and precious metals (16), whereas those rated low on both dimensions included wood (36) and wicker baskets (04). In contrast, stimuli such as bundles of cash (39) were rated as high in attractiveness but low in glossiness, whereas items such as metal cans (12) were rated as low in attractiveness but high in glossiness.

**Figure 2.**
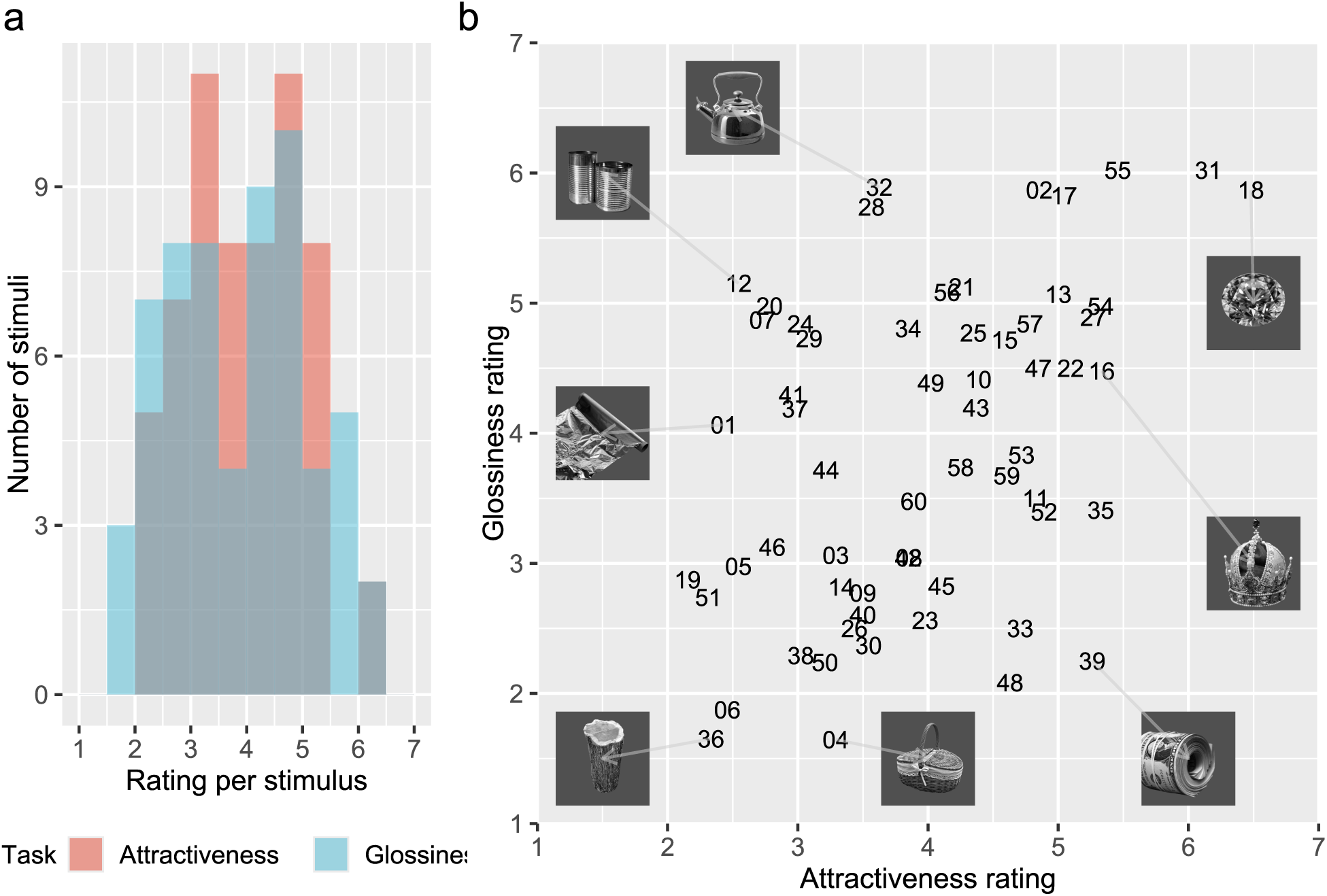
Ratings in the Experiment. (a) Histogram of the mean ratings for the 60 stimuli. The horizontal axis indicates the rating values, and the vertical axis indicates the frequency. The colors represent the rating tasks (red: attractiveness; blue: glossiness). (b) Scatterplot of the mean ratings for the 60 stimuli. The horizontal axis indicates attractiveness, and the vertical axis indicates glossiness. The numbers correspond to the stimulus IDs.

### 3.2 Pupil responses: Grand average

Figure 3 illustrates the average pupil responses for each rating level across the two tasks. Although the same set of stimuli was used, the pupil responses varied depending on the task. The peak of pupil constriction occurred approximately 1,000 ms after stimulus onset. Furthermore, apparent differences in mean pupil responses across rating levels were observed in both tasks. Specifically, in the attractiveness task, intermediate ratings were associated with greater pupil constriction, whereas in the glossiness task, higher ratings were associated with stronger constriction.

**Figure 3.**
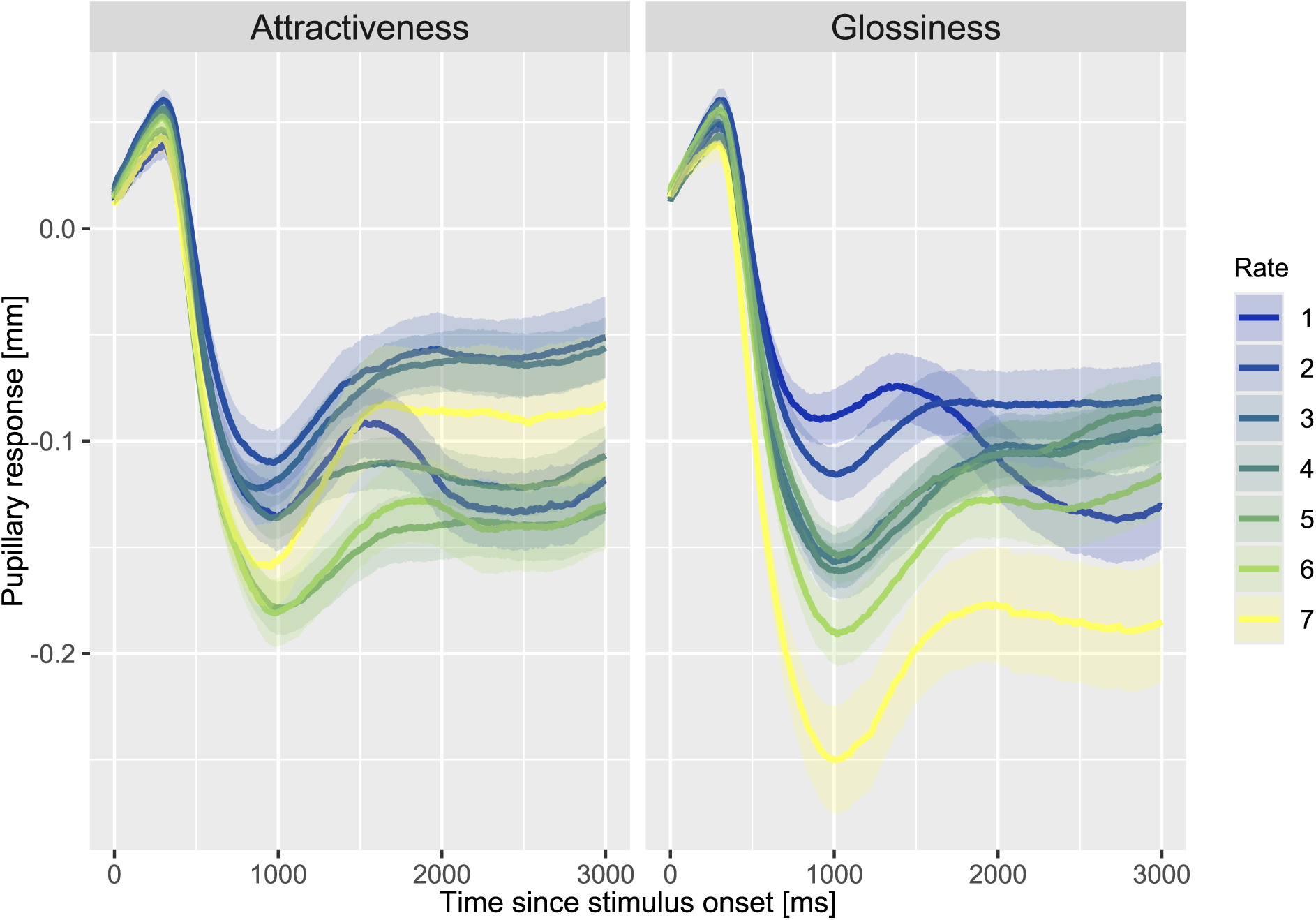
Average pupillary responses. The horizontal axis indicates the time from stimulus onset, and the vertical axis indicates the mean pupil response. Each color corresponds to a different rating level, and the shaded error bars represent the standard error of the mean.

### 3.3 Pupil responses: Temporal PCA

The temporal components that primarily accounted for the observed dynamics could not be identified on the basis of the average pupil responses across trials and rating levels. Therefore, to characterize the temporal dynamics of pupil responses, we applied tPCA to the raw pupil time series data, following established procedures (Scharf et al., 2022). The resulting factor loadings are shown in Figure 4. Since the first three components collectively accounted for 97.7% of the total variance, we limited our subsequent analyses and interpretations to these components. Factor 1 (72.5%) captured a late component characterized by a temporally broad and sustained response. Factor 2 (15.6%) reflected an earlier component with a more temporally confined profile, peaking approximately 1,000 ms after stimulus onset, consistent with the canonical pupillary light reflex. Factor 3 (9.6%) peaked even earlier and exhibited a transient profile, likely reflecting an orienting response associated with early sensory processing.

**Figure 4.**
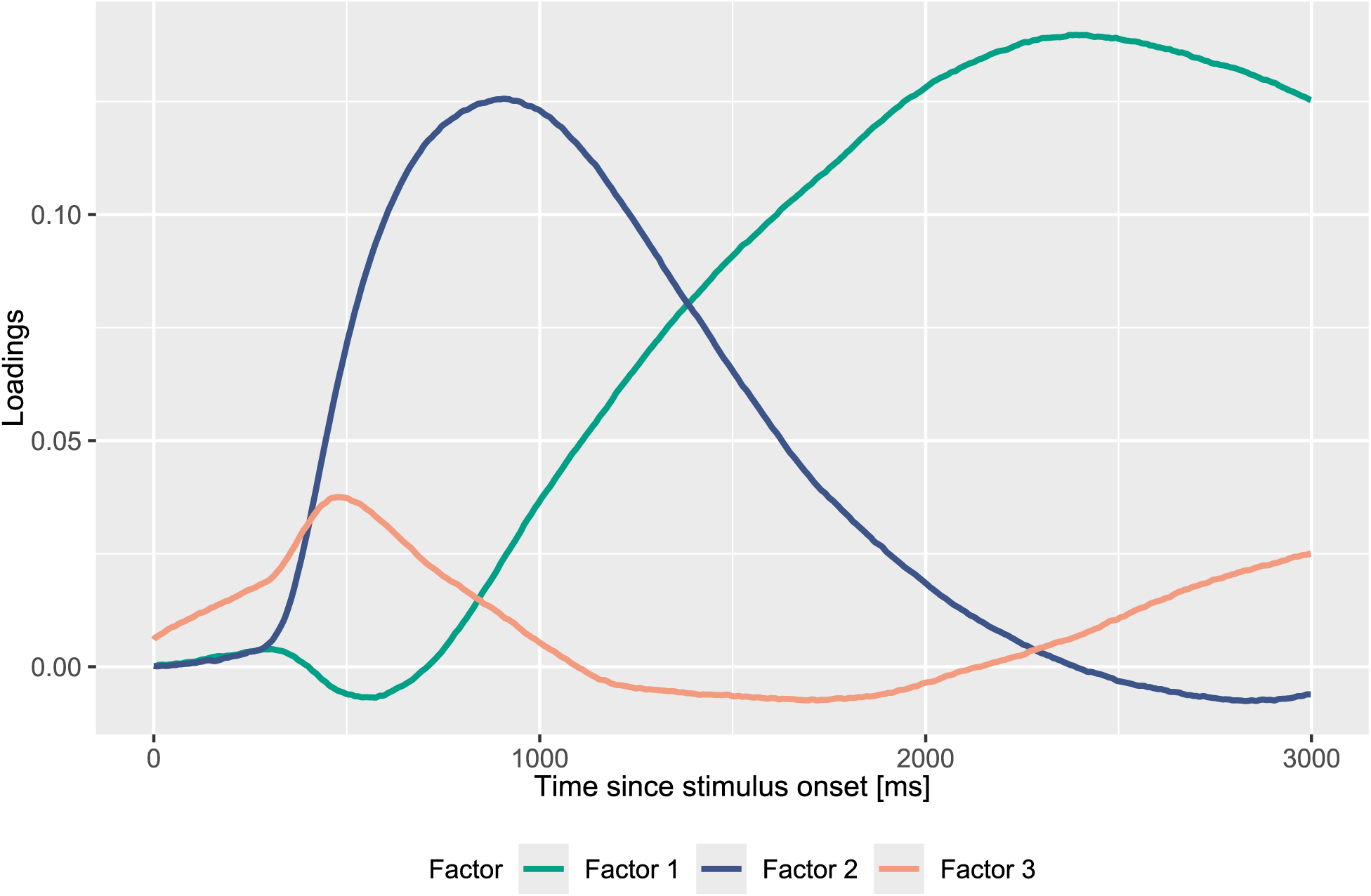
Temporal principal component analysis (tPCA) loadings. The horizontal axis indicates the time since stimulus onset, and the vertical axis indicates the factor loading. The three colored lines correspond to Factor 1 (green), Factor 2 (blue), and Factor 3 (orange).

Next, following the approach of a previous study (Wetzel et al., 2020), we computed single-trial component scores by multiplying each pupil response by the loading of each factor derived from the tPCA results. The results are presented in Figure 5, with colors indicating the rating task and subjective rating value.

**Figure 5.**
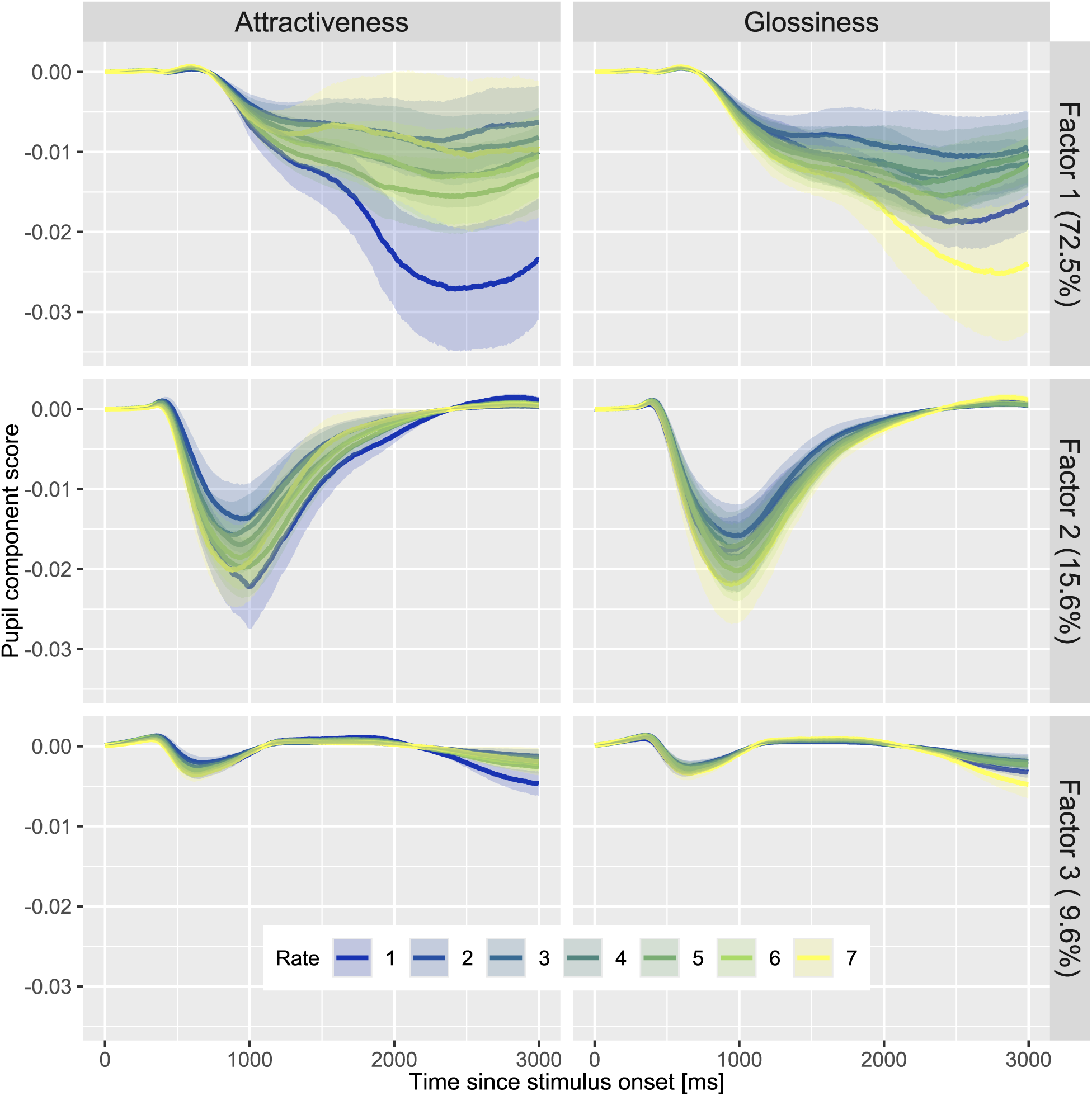
Pupil component scores derived from tPCA. The horizontal axis indicates the time since stimulus onset, and the vertical axis indicates the pupil component score. Each color corresponds to a different rating level, and the shaded error bars represent the standard error of the mean. The rows represent the three factors derived from tPCA, and the columns represent the two rating tasks.

As shown in Figure 5, the pupil responses associated with attractiveness were most prominently reflected in Factor 1. Specifically, delayed pupil constriction following the light reflex was observed in trials in which the participant provided a low attractiveness rating, whereas greater pupil dilation in the later phase was observed in trials in which the participant provided a high attractiveness rating.

In contrast, the pupil responses related to glossiness were captured in both Factor 1 and Factor 2. Regarding Factor 1, higher glossiness ratings were associated with stronger pupil constriction after the light reflex. Factor 2, which reflects the timing of the light reflex itself, was also associated with increased constriction with higher glossiness ratings. Notably, the magnitude of pupil constriction varied systematically with the level of perceived glossiness, indicating a parametric relationship.

### 3.4 Generalized additive model

To quantitatively assess whether pupil responses varied as a function of subjective ratings (Rate) and whether such variations differed across tasks, we applied GAM to the pupil component scores. Figure 6 presents the time course of partial effect differences between the attractiveness and glossiness tasks at fixed rating values of 1, 4, and 7 for Factor 1 (see also the top row in Figure 5).

**Figure 6.**
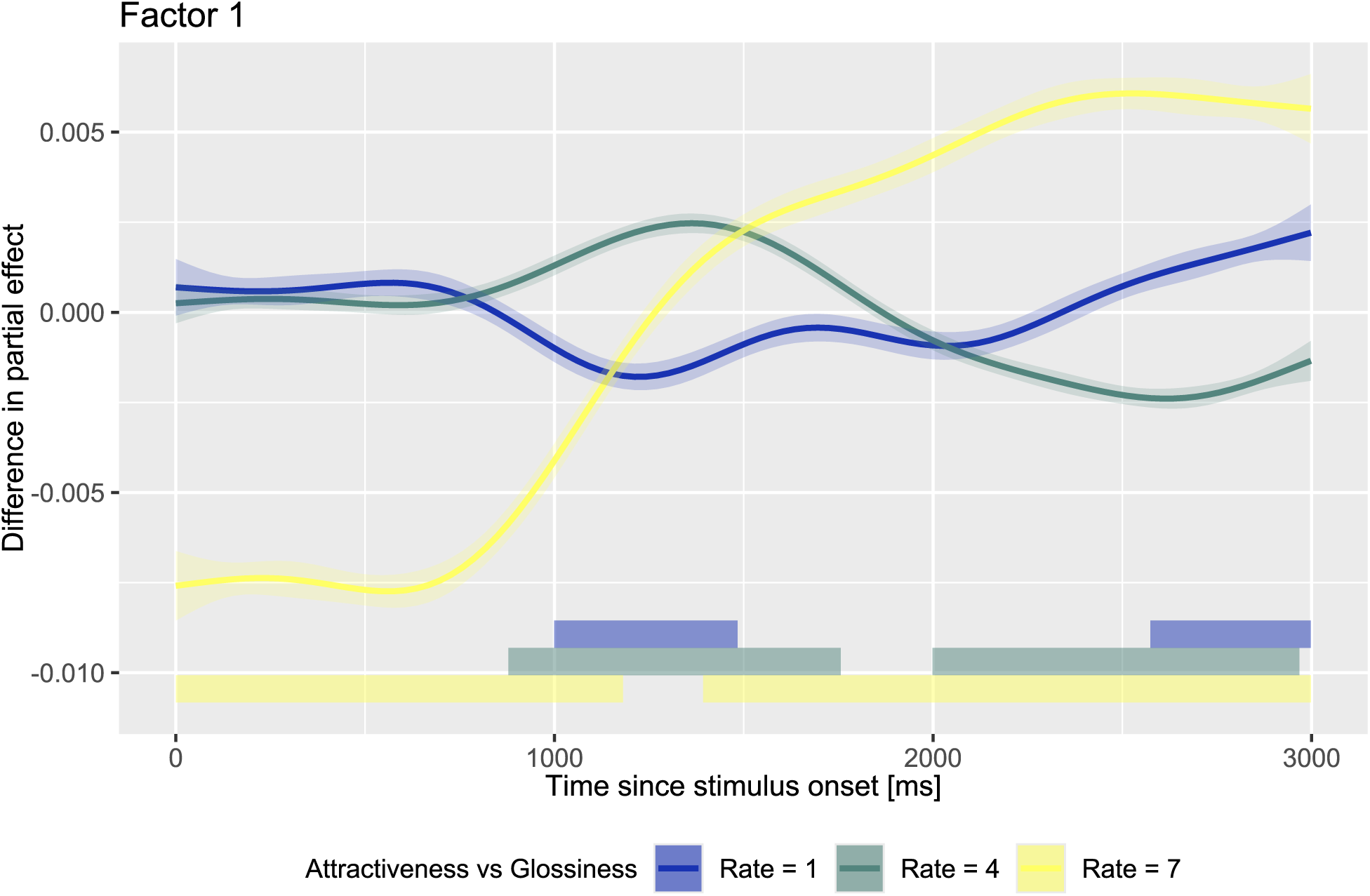
Results of the GAM analysis for Factor 1. The horizontal axis represents the time since stimulus onset, and the vertical axis indicates the difference in partial effects across conditions. Each line shows the difference between the attractiveness and glossiness tasks at each rating level (Rate). Positive values on the vertical axis indicate greater pupil dilation for attractiveness relative to glossiness or stronger pupil constriction for glossiness relative to attractiveness. The shaded areas represent 99% confidence intervals (p &lt; .01).

The analysis revealed pronounced task-related differences in the temporal dynamics of pupil responses, particularly at a rate of 7. Approximately 1200 ms after stimulus onset, the direction of the task effect appeared to reverse: before this point, the pupil diameter in the attractiveness task was smaller than that in the glossiness task (i.e., greater constriction), whereas after 1200 ms, it became larger (i.e., more dilation). At a rate of 7, statistically significant intervals were identified at 0–1181 ms and 1393–2998 ms, where the 99% confidence intervals of the estimated differences did not include zero (𝑝 < .01).

Although the differences at a rate of 1 were less pronounced, task-related effects were still evident during two temporal windows corresponding to the return phase of the pupillary light reflex (1000–1500 ms) and the late poststimulus period (2500–3000 ms). At a rate of 1, 999–1484 ms and 2574–2998 ms were identified as significant intervals (99% CI did not include zero, 𝑝 < .01). A similar pattern was observed at a rate of 4, with the task-related differences extending over a longer duration. The intervals 878–1756 ms and 1999–2968 ms were identified as significant (99% CI did not include zero, 𝑝 < .01).

The results of the GAM analysis for Factor 2 are shown in Figure 7. Within the interval corresponding to the pupillary light reflex (approximately 500–1600 ms after stimulus onset), characteristic task-related modulations were observed as a function of the rating. Compared with the glossiness task, pupil dilation was greater in the attractiveness task at a rate of 7, and this trend was also observed at a rate of 4. However, the opposite pattern—greater constriction in the attractiveness task—was observed at a rate of 1. In other words, stronger pupil constriction in the glossiness task was observed from a rate of 7 to a rate of 4. The significant intervals were wider and more pronounced than those observed for Factor 1 (Figure 6).

**Figure 7.**
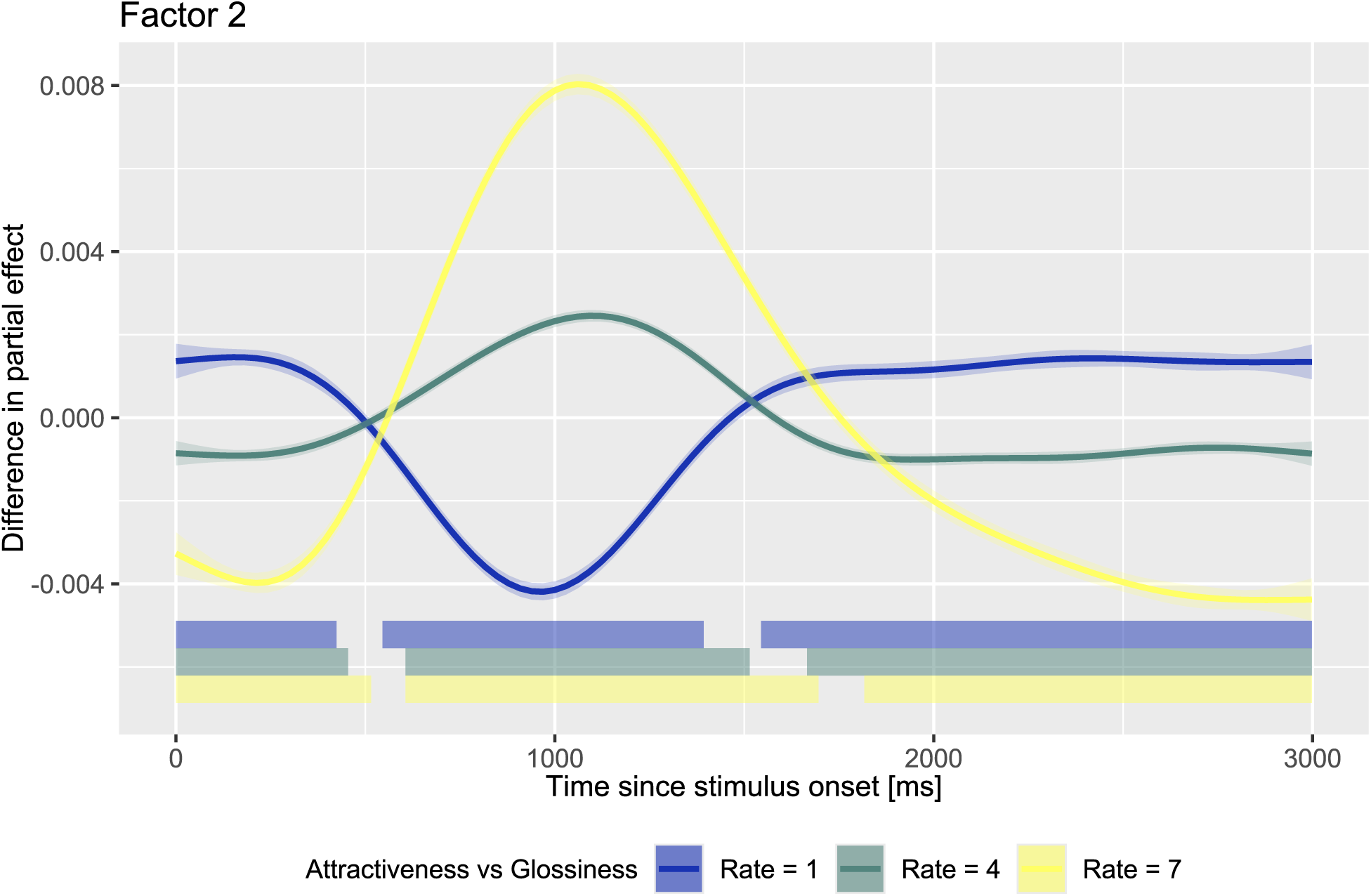
Results of the GAM analysis for Factor 2. The format was the same as that in Figure 6.

Similarly, the results of the GAM analysis for Factor 3 are shown in Figure 8. A distinctive pattern emerged at a rate of 7, where marked pupil dilation in the attractiveness task was observed beyond 2500 ms after stimulus onset. Notably, the absolute magnitude of the difference in the partial effect was smaller than that for Factors 1 and 2. In addition, the proportion of variance explained by Factor 3 was comparatively low (9.6%), further distinguishing it from the preceding factors. Taken together, although several intervals were identified as significant, caution is warranted when interpreting the task-related differences observed in this component.

**Figure 8.**
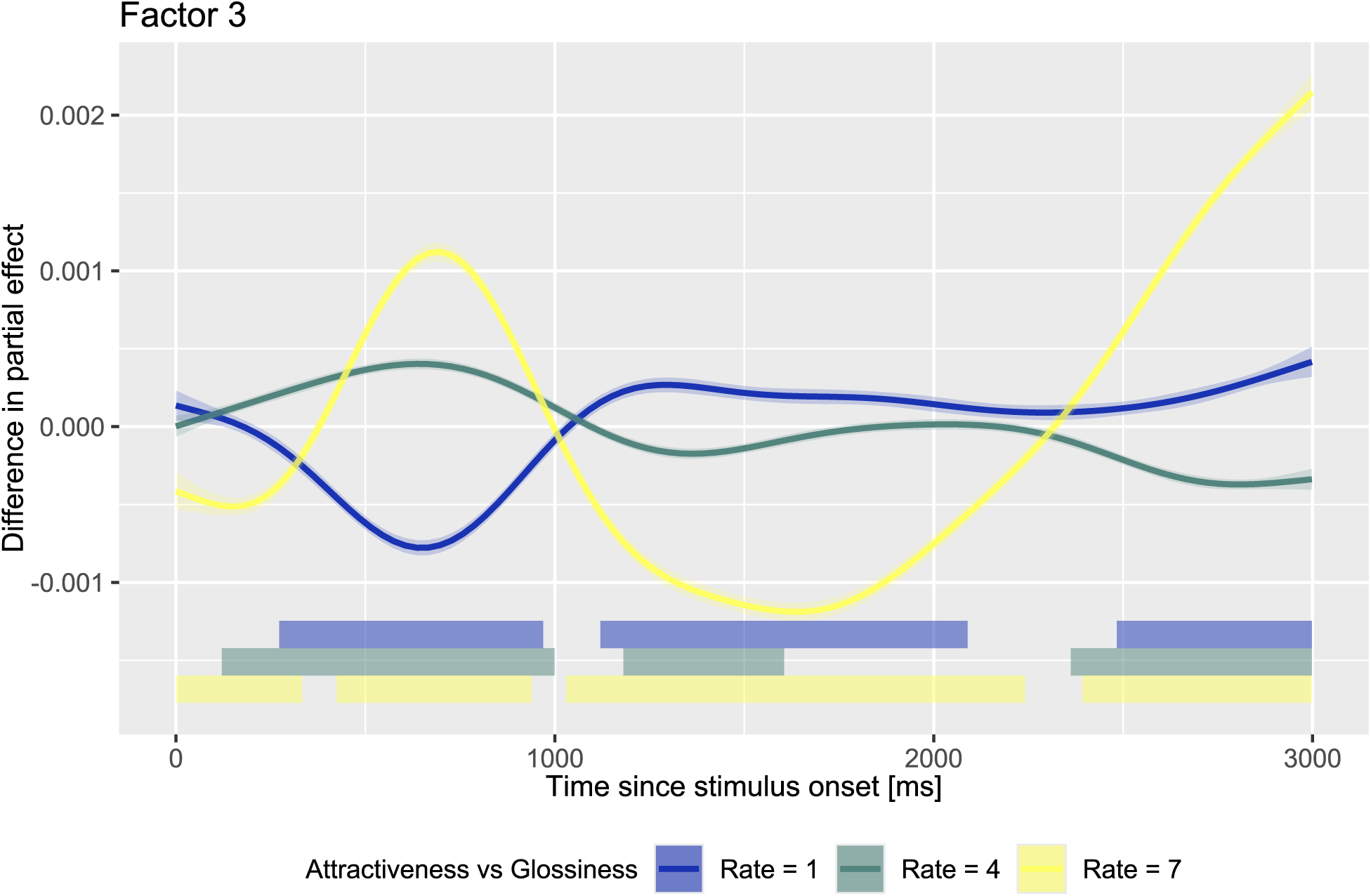
Results of the GAM analysis for Factor 3. The format was the same as that in Figure 6.

## 4. Discussion

In this study, we compared pupillary responses between two evaluative tasks —glossiness and attractiveness ratings—performed with identical visual stimuli. The rating data confirmed that participants perceived a broad range of glossiness and attractiveness, ranging from low to high values, indicating that the experimental manipulation was successful. Grand-average pupil traces revealed a common pattern of light-induced constriction followed by a gradual return to baseline; however, the amplitude and temporal characteristics of this response varied as a function of task and subjective rating. tPCA revealed distinct pupil response components that differed systematically across factors, tasks, and rating levels. Furthermore, GAMs of these components revealed consistent task-dependent differences in pupil dynamics. Taken together, these findings demonstrate that pupillary responses are modulated not only by the physical properties of the stimulus but also by the nature of the evaluative task and the subjective rating assigned to the same visual input.

Few behavioral studies have concurrently assessed both physical and affective texture qualities—such as glossiness and attractiveness—using the same set of visual stimuli, as illustrated in Figure 2a. One notable exception is the study by Fleming et al. (Fleming et al., 2013), in which participants were asked to rate texture images of everyday objects using various descriptive adjectives. They reported that interparticipant correlations were relatively high (approximately 0.6–0.8) for physical material attributes (e.g., glossiness, transparency) but substantially lower (approximately 0.3) for more subjective impressions such as prettiness. Consistent with this trend, the present study revealed interparticipant correlations of 0.53 for glossiness (a physical texture quality) and 0.41 for attractiveness (an affective texture quality). These results suggest that physical texture impressions tend to be more universally perceived and are more closely related to the optical properties of the object itself, whereas affective impressions such as attractiveness are more susceptible to individual differences and subjective interpretation.

We successfully replicated the main finding of Tamura et al. (2024)—namely, a systematic relationship between subjective ratings of material quality and pupil responses to everyday objects—with the overall average pupil data (Figure 3) (Tamura et al., 2024). While their analysis focused primarily on summarizing pupillary responses using a single index, namely, the magnitude of maximum pupil constriction, we extends this approach by applying time-resolved methods such as tPCA and GAMs. This analytic advancement represents a novel contribution, as we examined the full temporal dynamics of the pupil response without relying solely on peak amplitude or latency measures. The tPCA and GAM results revealed that even when the same set of visual stimuli was used across conditions, the temporal profile of the pupillary response systematically varied as a function of the evaluative task.

As illustrated in Figure 5, each factor appears to capture temporally distinct components of the pupillary response: Factor 1 (F1) corresponds to a late dilation-related component, Factor 2 (F2) reflects an earlier constriction component likely driven by the pupillary light reflex, and Factor 3 (F3) represents an initial orienting response. Notably, F1 was associated with greater dilation at higher attractiveness ratings, suggesting that more attractive stimuli elicit heightened attention (Binda et al., 2013a; Bombeke et al., 2016; Mathôt et al., 2013; Naber et al., 2013) and increased arousal (Bradley et al., 2008; Kuraguchi & Kanari, 2020, 2021; Laeng et al., 2013; Oliva & Anikin, 2018), which is consistent with prior findings. While pupil dilation has frequently been associated with high attractiveness, Liao et al. (2021) reported the opposite effect, namely, transient pupil constriction in response to highly attractive faces (Liao et al., 2021). Interestingly, a similar pattern of early constriction emerged in the present study within the early time window (500–1500 ms) of both F1 and F2, as revealed by the GAM analyses (Figures 6 and 7). This result is particularly notable given the correspondence in timing and response polarity. However, several methodological differences limit direct comparisons: facial stimuli were used in Liao et al.’s study, glossiness was not included as a comparison task, and unseparated pupil signals were analyzed rather than decomposed components. In contrast, we applied tPCA to general object images and systematically compared glossiness and attractiveness evaluations among the same participants. Despite these differences, our findings suggest that automatic pupillary modulations may emerge early in response to visually attractive stimuli, even before conscious evaluations are completed. Moreover, consistent with Santos et al. (2023), our results demonstrate that physiological responses can vary depending on evaluative instructions, even when the visual input remains constant (Santos et al., 2023). Importantly, while Santos et al. manipulated the evaluative context between participants, our within-subjects design allowed for a more direct comparison of pupil dynamics across tasks using identical stimuli.

In addition to the effects observed for F1, clear task-dependent differences in pupillary responses were observed in the analysis of the F2 component (Figure 7). Specifically, our results replicated previous findings showing that higher glossiness ratings were associated with stronger pupil constriction (Figures 3 and 5), which is consistent with the findings of Tamura et al. (2024) (Tamura et al., 2024). At the peak of the light-induced pupillary constriction—approximately 1000 ms after stimulus onset—the difference in pupil diameter between the lowest (rating 1) and highest (rating 7) glossiness levels reached approximately 0.15 mm. This value is comparable to, or even exceeds, the 0.1 mm effect size reported in a prior study (Tamura et al., 2024) (see the right panel in Figure 3). The magnitude of this modulation is also consistent with previous studies reporting pupil constriction in response to illusory brightness or high-level image content that alters subjective luminance perception (Binda et al., 2013b; Laeng & Endestad, 2012; Naber & Nakayama, 2013).

Furthermore, task-related differences in pupil responses were observed for F3. Given that the observed modulation occurred earlier than the pupillary constriction typically associated with the light reflex, it likely reflects a transient response associated with novelty or orienting mechanisms. As shown in Figure 5 (see also Figure S1 for the zoomed-in view), pupil changes appear to be linked to attractiveness ratings, suggesting that the attractiveness judgment task—due to its greater emotional and social salience —may have induced the differential allocation of early attentional resources via the superior colliculus (Wang et al., 2014; Wang & Munoz, 2015). Nevertheless, considering that the variance explained by this component was comparatively low (9.6%), this finding should be interpreted cautiously.

In the present study, we observed systematic differences in the temporal profile of pupillary responses depending on the evaluative task, despite the use of physically identical stimuli. The tPCA results revealed distinct latent components across tasks (attractiveness vs. glossiness), suggesting that pupil dynamics are not strictly constrained by a fixed physiological manifold (Blini et al., 2024) but can instead exhibit plasticity depending on the cognitive context imposed by the task. This finding differs from the results of Blini et al. (2024), who reported that pupillary responses to numerical stimuli followed a shared low-dimensional trajectory across different evaluative tasks, indicating a robust physiological structure that was largely invariant to cognitive framing. However, it is important to note a key methodological distinction between the two studies in terms of how “stimulus constancy” was operationalized. In Blini et al.’s study, stimulus properties such as numerical value and brightness varied across tasks, whereas we employed strictly identical physical stimuli across conditions, allowing for a more controlled comparison.

Therefore, our findings suggest that pupil dynamics are more flexibly modulated by task context than previously assumed and offer a complementary and potentially expansive perspective on the notion of a fixed pupillary manifold. Specifically, our results suggest that the possibility that the temporal structure of pupillary responses is shaped not only by the intrinsic properties of the stimulus but also by the evaluative process itself.

We controlled the luminance levels of all the stimuli to isolate the psychosensory components of the pupillary response—those arising above and beyond the luminance-driven light reflex—across different evaluative tasks. Recent studies have reported that pupillary responses can exhibit orientation-dependent modulations even when luminance is held constant (Parrella et al., 2025). This suggests that despite controlling for overall luminance, subtle differences in the spatial configuration of the stimulus—such as whether the object occupied a vertically or horizontally elongated region —may still influence the pupillary response. Thus, care must be taken in interpreting psychosensory effects, as spatial anisotropy in the stimulus layout could contribute to task-dependent pupil dynamics.

Another possible explanation for the observed differences in pupil responses between the attractiveness and glossiness tasks is that each evaluation was conducted in a separate block. Performing a series of judgments within the same task block may have induced a specific evaluative mode or mental set unique to each texture dimension, which could have influenced the pupillary responses. Although the order of task blocks was counterbalanced across participants, this measure does not eliminate the possibility that block-level task context contributed to the observed effects. This methodological consideration highlights the value of alternative paradigms—such as interleaving trials of different evaluative tasks within a single block—to minimize task-specific preparatory states and better isolate the cognitive mechanisms underlying pupillary modulations.

While the observed latency differences in pupil responses—particularly the finding that attractiveness-related modulations emerged later than glossiness-related modulations —can be interpreted considering the hypothesis that surface property evaluations are hierarchically organized (Komatsu & Goda, 2018), we do not claim to directly test the underlying neural framework. Instead, our findings provide indirect physiological evidence that low-level and high-level material qualities may engage temporally distinct processes. Nevertheless, pupil diameter constitutes a one-dimensional physiological index. Although this measure is informative, it offers limited insight into the spatiotemporal dynamics of the neural mechanisms involved. Future research should incorporate multimodal approaches such as electroencephalography or functional neuroimaging to better resolve the stages of visual and affective evaluation, particularly in relation to surface perception.

Moreover, we focused exclusively on glossiness and attractiveness as representative examples of low- and high-level material attributes. Whether similar patterns of pupillary modulation extend to other perceptual dimensions (e.g., roughness, transparency) or material categories (e.g., fabric, metal) remains an open question. Investigating pupillary responses across a broader range of attributes is essential for testing the generalizability of the present findings and refining theoretical models of material perception.

## 5. Conclusion

In this study, we examined how pupillary responses differed when participants evaluated the same set of stimuli in terms of glossiness and attractiveness. tPCA and GAMs revealed that higher glossiness ratings were associated with greater pupil constriction at an earlier, light-reflex–like timing. In contrast, higher attractiveness ratings were linked to greater pupil dilation at a later stage. These results may reflect differences in the temporal stages of visual processing recruited for each evaluation task. Overall, our findings highlight that task-dependent pupillary dynamics can serve as a promising physiological index for investigating hierarchical processing in material perception, with potential applications in cognitive neuroscience and affective engineering.

