## Supplementary Information for "Task-dependent pupillary responses to glossiness and attractiveness judgments"

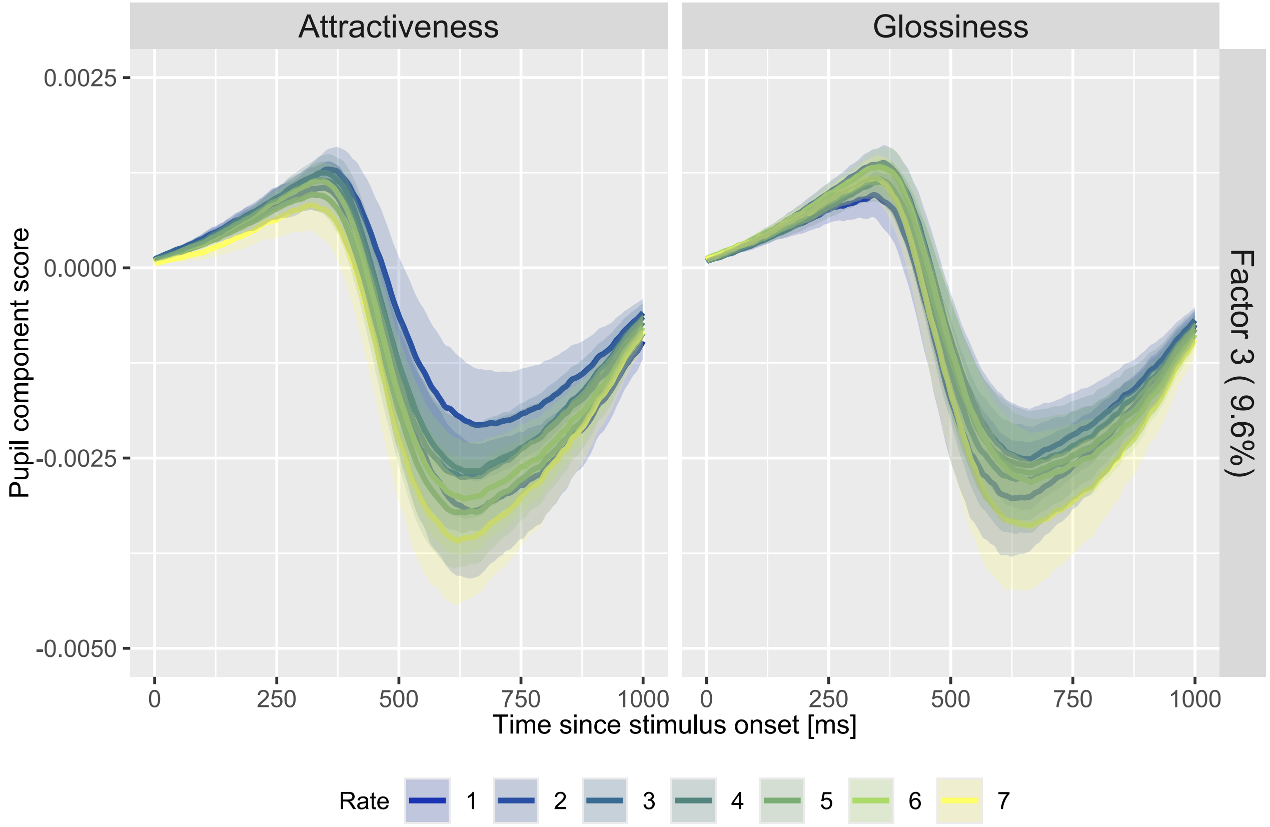


**Figure S1.** Zoomed-in view of the pupil component scores for Factor 3 derived from tPCA, focusing on the 0–1000 ms interval after stimulus onset. The plotting format is identical to that of Figure 5. Please refer to the caption of Figure 5 for axis labels, color coding, and layout details.
